# Dendritic sodium spikes endow neurons with inverse firing rate response to correlated synaptic activity

**DOI:** 10.1101/137984

**Authors:** Tomasz Górski, Romain Veltz, Mathieu Galtier, Hélissande Fragnaud, Jennifer S. Goldman, Bartosz Teleńczuk, Alain Destexhe

**Affiliations:** Unité de Neurosciences, Information et Complexité, Centre National de la Recherche Scientifique, Gif-sur-Yvette, France; European Institute for Theoretical Neuroscience Paris, France; Inria, Sophia Antipolis, France

**Keywords:** Dendritic integration, Synaptic input correlations

## Abstract

Many neurons possess dendrites enriched with sodium channels and are capable of generating action potentials. However, the role of dendritic sodium spikes remain unclear. Here, we study computational models of neurons to investigate the functional effects of dendritic spikes. In agreement with previous studies, we found that point neurons or neurons with passive dendrites increase their somatic firing rate in response to the correlation of synaptic bombardment for a wide range of input conditions, i.e. input firing rates, synaptic conductances or refractory periods. However, neurons with active dendrites show the opposite behavior: for a wide range of conditions the firing rate decreases as a function of correlation. We found this property in three types of models of dendritic excitability: a Hodgkin-Huxley model of dendritic spikes, a model with integrate-and-fire dendrites, and a discrete-state dendritic model. We conclude that neurons equipped with with fast dendritic spikes confer much broader computational properties to neurons, sometimes opposite to that of point neurons.

## 1 Introduction

Increasing evidence shows that nonlinear integration of synaptic inputs in dendrites is crucial for the computational properties of neurons. A major role in the integration is played by dendritic spikes: regenerative currents through Na^+^, Ca^2+^ or NMDAr channels. The first evidence of dendritic spikes came from field recordings [1–5], corroborated by the intracellular recordings [6–8]. The repertoire of techniques was further enlarged by patch clamp [9–13] and optical methods. Calcium imaging allowed for the direct observation of calcium spikes [14–18], and glutamate uncaging and voltage sensitive dyes led to the discovery of NMDA spikes [19,20].

Dendritic spikes allow for more subtle integration of synaptic input than in a passive dendrite. A single dendritic branch can act as a coincidence detector, generating a spike when exposed to synchronized input [20–22]. The propagation of dendritic spikes generated in the distal part of dendritic tree can be gated by synaptic input in the proximal region, as was shown for hippocampal CA1 pyramidal neurons [23], and L5 pyramidal neurons [24]. After the initiation of a dendritic spike, sodium channels inactivate and the branch switches into a refractory state which crucially affects integration [25]. Backpropagating action potentials also play an essential role in spike time-dependent plasticity [26–28], and the participation of local dendritic spikes has been implicated in long-term potentiation [29, 30].

Sodium spikes can propagate in many cell types: neocortical pyramidal cells [9, 31–34], hippocampal CA1 and CA3 pyramidal cells [35–39], interneurons [40] or thalamic neurons [41]. Models predicted that, *in vivo*, the presence of synaptic background activity should greatly enhance the initiation and propagation of dendritic spikes [42, 43]. This suggests that there can be a heavy traffic of dendritic spikes, as indeed found in recent dendritic recordings in awake animals [44]. Therefore interactions between dendritic spikes likely play an important role in dendritic integration.

These interactions can be conveyed by the refractory period which follows each spike [45], and can prevent initiations of subsequent spikes, or can result in a collision and annihilation of spikes [46]. Here, comparing different types of computational models, we show that interactions between dendritic spikes can change the way the correlation of synaptic input affects the firing rate of the cell. For single-compartment neurons and neurons with passive dendrites an increase of correlation of synaptic input is known to cause an increase of firing rate [47–50], while this relation can be reversed only for high input intensities. We show that neurons with active dendrites behave the opposite way: for wide range of input conditions, the firing rate varies inversely proportionally to the level of correlation. We discuss possible consequences of this property at the network level.

## 2 Results

We first show the phenomenon of inverse correlation processing using a multi-compartment Hodgkin-Huxley model with active dendrites. Next, we investigate the impact of the refractory period duration on inverse processing using a multi-compartment integrate-and-fire model. Finally, we show that this phenomenon is also present in simplified discrete-state dendritic models.

### 2.1 Model of correlated synaptic activity

In all models studied here, we considered neurons subject to *in vivo*–like synaptic activity. In particular, we aimed at investigating the effects of synaptic noise correlations on the firing of the cell. The synapses were located on the somatic and dendritic compartments and were of two types: excitatory AMPA synapses (with reversal potential *E_e_* = 0 mV) and inhibitory GABA_A_ synapses (*E_i_* = −75 mV), with five times more excitatory than inhibitory synapses [51].

We considered the case where the pre-synaptic spikes triggering the excitatory inputs are correlated. There might be two biological sources of such correlation. First of all, in cortical networks neurons are known to share some of their synaptic inputs which inevitably makes their spikes correlated. Since this type of correlation is common to larger populations of neurons, we call it the global correlation [52]. In addition, a single axon can create several synapses on close-by segments of the dendrite activating them at the same time [53], so that the correlation of the input spikes within the same dendritic segments (local correlation) can be higher than the global correlation.

To generate such locally and globally correlated synaptic inputs, we used a previously proposed algorithm [54]. From a global spike train *G* obtained from Poisson process, we draw random spikes with probability *r_G_*. The spike train *L_k_* thus obtained corresponds to all synapses on a single dendritic compartment. Finally, the spike train 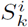 for a synapse *i* situated on compartment *k* is built by drawing spikes from *L_k_* with local probability *r_L_* [Fig. 1].

**Fig. 1.**
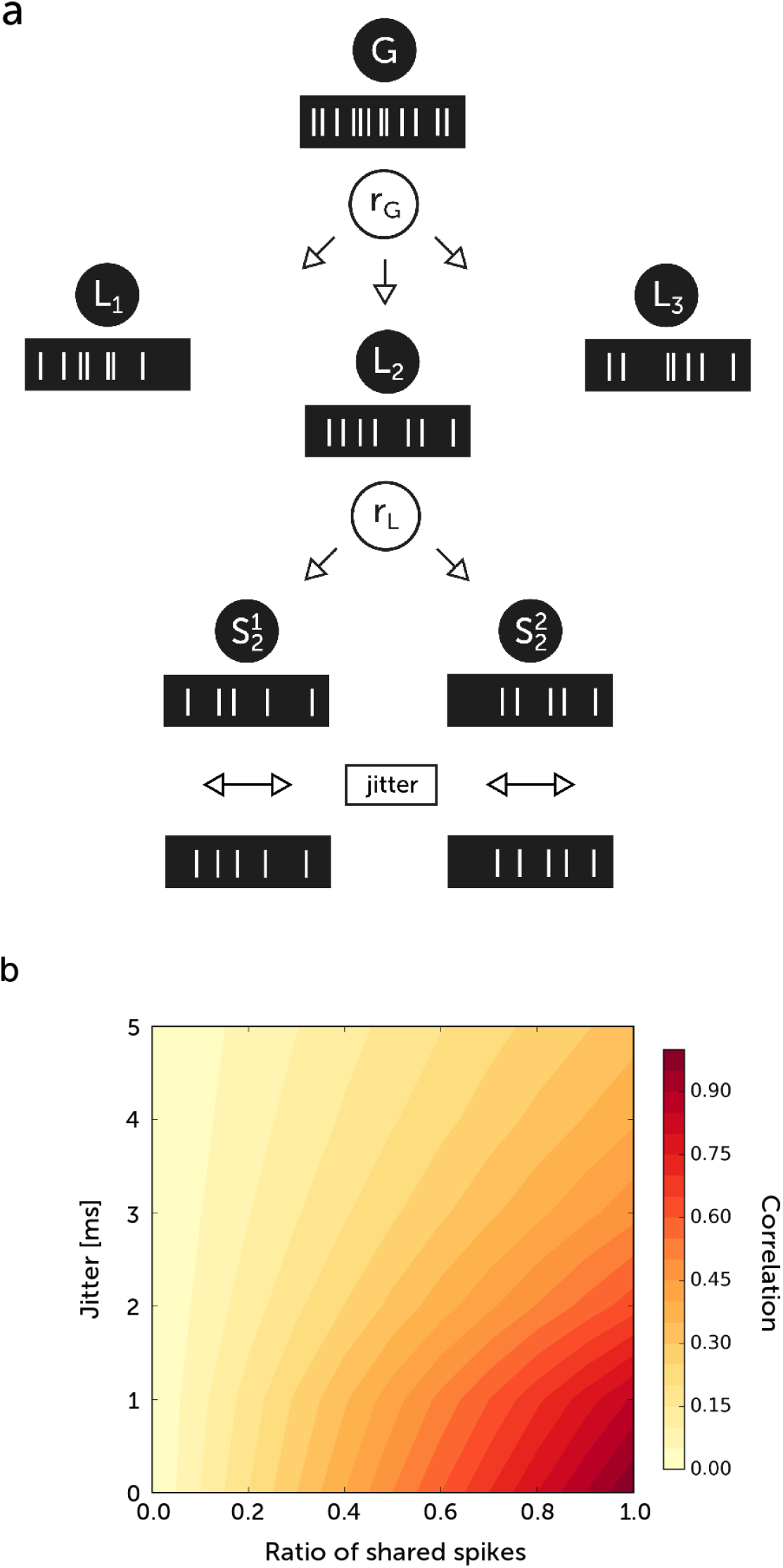
Model of correlated synaptic activity. (a) From the global spike train *G* spike times are distributed with probability *r_G_* to the local spike trains *L_k_* corresponding to compartments. From the local spike trains spikes are distributed with probability *r_L_* to synapses 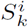. The spike times are then shifted in time with random jitter. (b) The dependence of correlation measure 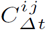 between two spike trains *i* and *j* on a ratio of shared spikes and on the jitter. To calculate the correlation the time range of cross-covariance function integral was *∆t* = 2 ms. The mean rate of the process was 20 Hz, and its duration was 50 s.

In this model, the ratio of shared spikes between synapses situated on the same compartment is equal to *c_L_* = *r_L_* and the ratio of spikes shared by synapses situated on different compartments is equal to *c_G_* = *r_L_r_G_*. Note that *c_L_* is always greater than or equal to *c_G_*. At the end, we added a jitter to each spike time to obtain desynchronized input. The jitter is drawn from exponential distribution with a standard deviation *τ_j_* with equal probability for positive and negative values. To obtain the time-dependent synaptic conductances, we convolve the resulting spike trains with an exponential function reflecting the change of synaptic conductance due to a single spike.

### 2.2 Response to correlated synaptic activity in biophysical dendritic models

We first apply this model of correlation processing to Hodgkin-Huxley model of neuron with a dendrite (see Methods). In our model the dendritic spikes can be one order of magnitude more frequent than somatic spikes (Fig. 2), value similar to that observed in vivo [44]. In Fig. 3, we show how dendritic sodium spikes propagate and collide in the model. We investigate how interactions between dendritic spikes can affect the response of a neuron to correlated synaptic activity. We run simulations of the model under correlated synaptic activity. For each run, we measured the firing rate of somatic spikes and asked how this quantity is affected by the ratio of shared spikes (Fig.4). For each ratio we performed 20 runs with a duration of 20 s. With increasing density of active channels in the dendrite, the somatic firing response changes to become *inverse*, i.e. we observe a decrease of somatic firing rate with the correlation of synaptic input (Fig.4a). To adequately compare the multicompartment neuron with a point neuron, we have scaled the synaptic conductances in the point neuron to emulate filtering of EPSPs by the dendrite. The inverse response of a neuron with dendrite can be enhanced by the increase of input firing rate and the increase of synaptic weights (Fig.4a,b). In contrast, in the point neuron model, the firing rate response generally increased with correlation, as found previously [47, 48]. It could also remain approximately constant for high values of the synaptic input (> 40 Hz). The increase of synaptic weights resulted in a more pronounced inverse response to correlation of synaptic input. The reason for this effect is that, for low weights, more synaptic potentials remain sub-threshold and summed synaptic inputs from single or multiple compartments are necessary to trigger a single spike. Hence, a higher input correlation leads to an increase in the probability of generating a spike. On the other hand, for larger synaptic weights, few synaptic inputs may trigger a spike, whose propagation to the soma is limited by the effects of the refractory period and collisions. In this regime, a higher correlation means higher probability of stopping spike propagation by the refractory periods of preceding spikes. Therefore, we observe a stronger decrease in the somatic firing rate.

**Fig. 2.**
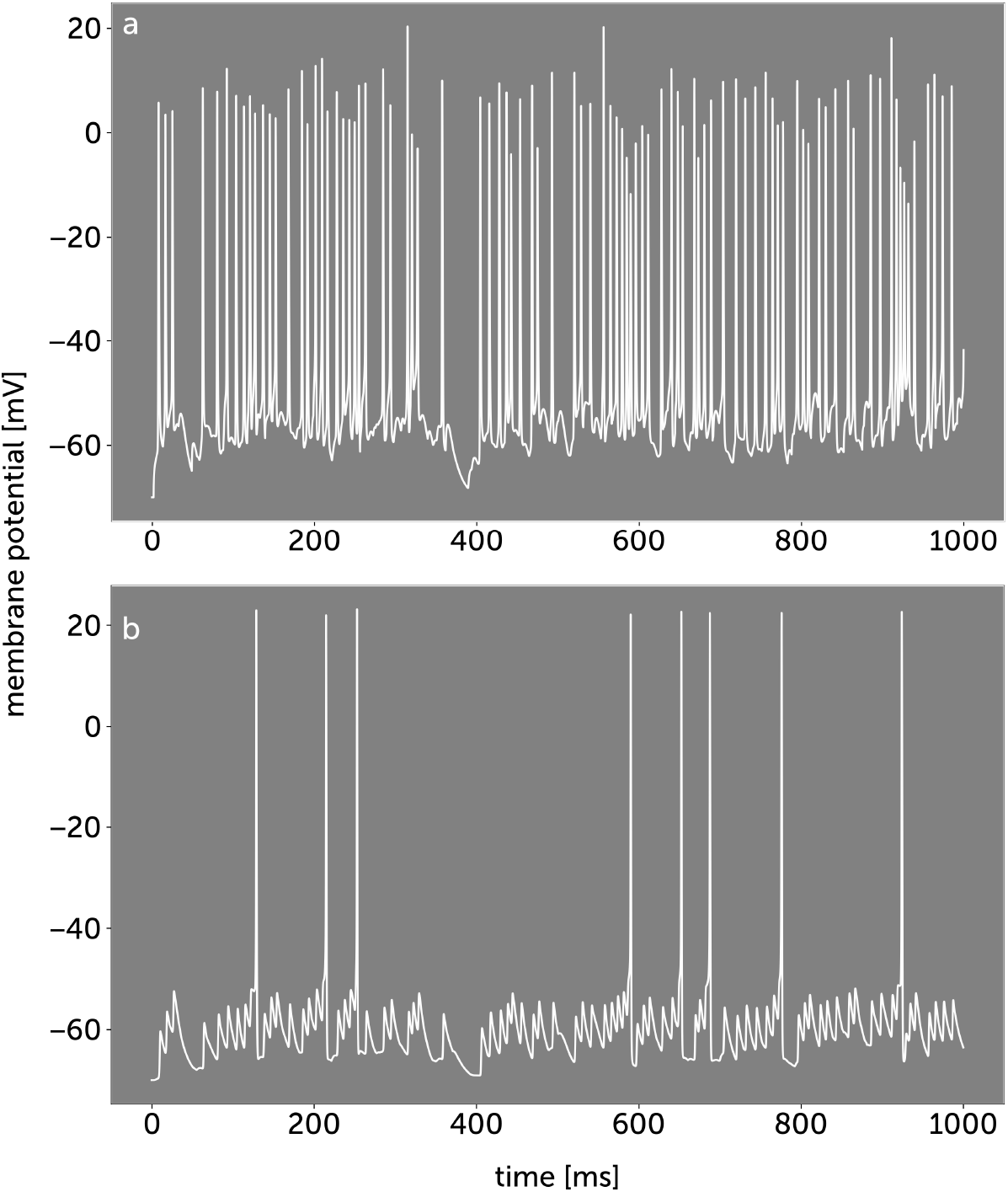
Comparison of dendritic and somatic spikes. (a) Membrane voltage in the dendrite 500 *μ*m from soma. (b) Membrane voltage in soma. Most of the dendritic spikes cannot actively invade soma causing depolarizations with amplitudes of few millivolts.

**Fig. 3.**
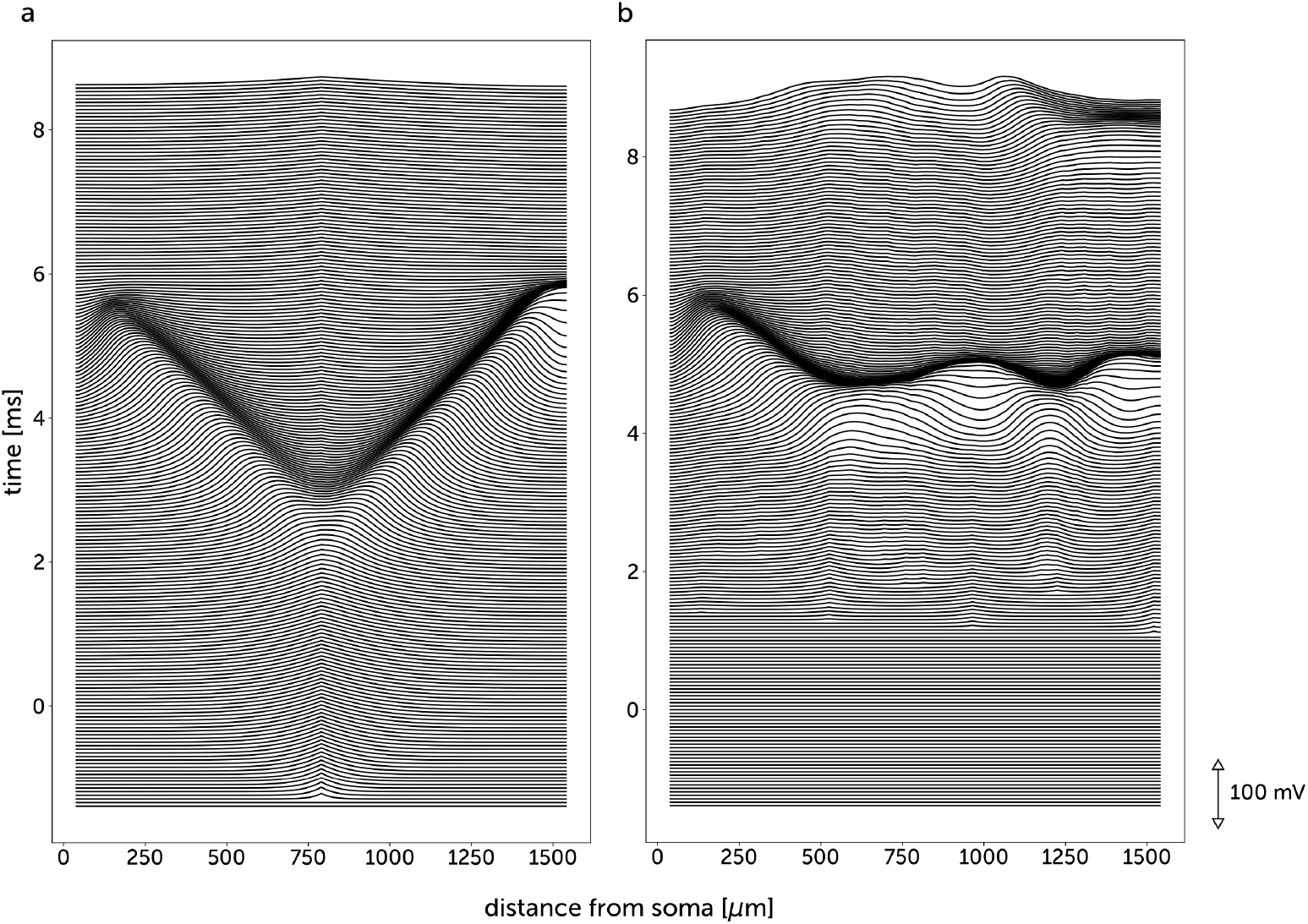
Propagation and collision of dendritic spikes in a Hodgkin-Huxley model. Each line represents the profile of membrane voltage at a given instant. (a) A sodium spike is created near the center of the dendrite and propagates toward the soma and the distal dendritic end. (b) The correlated synaptic bombardment (*c_G_* = 0.1, *τ_j_* = 5 ms) creates two sodium spikes (around 600 *μ*m and 1200 *μ*m from soma) which wavefronts collide and cancel due to the refractory period.

**Fig. 4.**
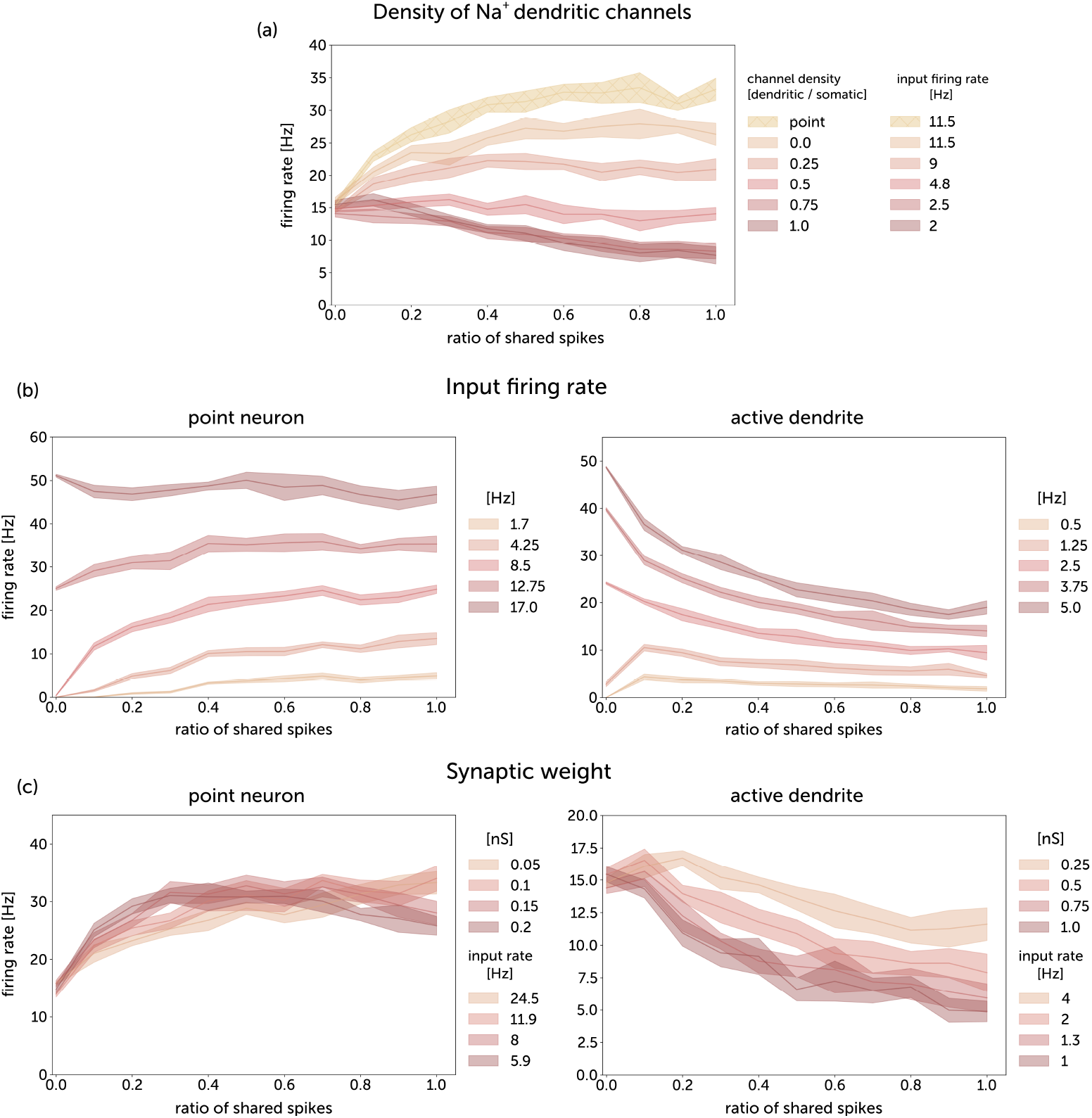
Somatic firing rate responses to correlated synaptic activity for a Hodgkin-Huxley point neuron and for a Hodgkin-Huxley neuron with dendrite. (a) Firing responses for different densities of sodium channels in a dendrite. While changing dendritic densities, the somatic densities were left unchanged (12 mS*/*cm^2^ for Na^+^ conductance and 7 mS/cm^2^ for K^+^ conductance. The responses of the neuron with dendrite are compared with the responses of the point neuron with scaled synapses. The synaptic conductances were 0.5 nS for the neuron with dendrite, and 0.105 nS for the point neuron. (b) Firing responses for different input firing rates for a Hodgkin-Huxley point neuron with scaled synapses and for a Hodgkin-Huxley neuron with dendrite. (c) Firing responses for different synaptic conductances for a point neuron and a neuron with dendrite. *For plots (b) and (c):* the dendritic Na^+^ channel density of the multi-compartment neuron was equal to the somatic channel density (12 mS*/*cm^2^). *For all plots:* in each of the 200 dendritic compartments of the neuron with dendrite, there was one excitatory synapse; 40 inhibitory synapses were located on the soma. For the point neuron, all 200 excitatory synapses were placed on the soma. The ratio of shared spikes for the multicompartment neuron is therefore c_G_. The time constant of jitter for the generation of correlated input spike trains, *τ_j_* was 10 ms.

To investigate the mechanism of inverse processing, we measured the frequency of dendritic spikes simultaneously with the frequency of somatic spikes. With an increase of the density of dendritic sodium channels, the ratio of dendritic to somatic spikes increases abruptly, reaching values around 10 for the same density of somatic and dendritic channels (Fig. 5a). With an increase of synaptic input correlation, the number of dendritic spikes decreases due to spike cancellations, which results in a weaker depolarization of the soma and a lower somatic firing rate (Fig. 5b.)

**Fig. 5.**
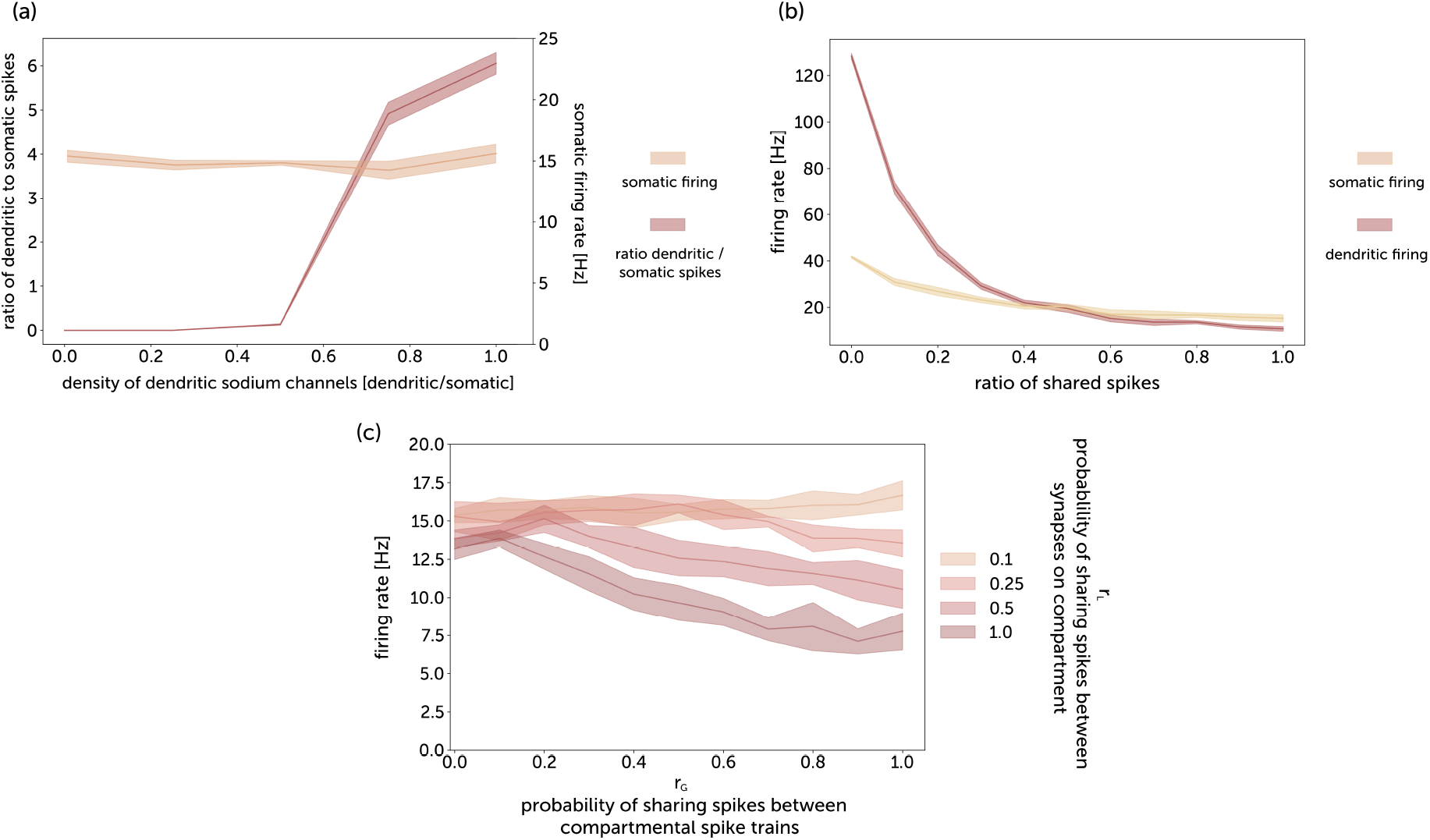
Somatic and dendritic spikes. (a) Ratio of dendritic spike firing to somatic spike firing as a function of the density of dendritic sodium channels. The somatic firing rate was kept at a nearly constant level of 15 Hz by adjusting the input firing rate while changing channel densities. The input spike trains were uncorrelated. The dendritic spikes were detected in the middle of the dendrite 500 *μm* from the soma. (b) Somatic and dendritic firing responses to correlated synaptic activity. (c) Somatic firing response on global correlation r_G_ for different values of local correlation r_L_. The dendrite consisted of 50 compartments of size 20 *μm*, with 4 excitatory synapses on each compartment yielding the same total number of synapses as for the previous simulations. *For all plots*: The synaptic conductances were 0.5 nS. The time constant of jitter for the generation of correlated input spike trains, *τ_j_* was 10 ms.

We checked how local correlations affect the firing rate response (Fig. 5c). We observed that neurons with active dendrites display a stronger inverse response to global correlations for more locally synchronized synaptic activity. Hence increasing local correlations acts as increasing effective synaptic weights (cf. Fig. 4c). This interplay between local and global correlation can largely affect the properties of dendritic integration.

To investigate the impact of the duration of refractory period on inverse processing, we considered an integrate-and-fire type model. In particular, we used a multicompartment exponential integrate and fire model, which can be extended to include the neuronal adaptation [55]. We extended the exponential integrate and fire model with axial currents and spike waveform to adapt it for modeling multi-compartment dendrites (see Methods for the equations of the multi-compartment model). With this modification, the model can produce a spike propagating across the tapered linear dendrite, but the propagation can be stopped by the tunable refractory period caused by other spikes.

The morphology of the integrate and fire model was the same as for the Hodgkin-Huxley model (see Methods). In the same way as for Hodgkin-Huxley model, we measure the firing rate of somatic spikes as a function of the ratio of shared spikes.

For the neuron with dendrite, the increase of the duration of the refractory period results in a stronger inverse response to correlation (Fig. 6). While for the point neuron model, the response remained positive even for 10 ms refractory period.

**Fig.6.**
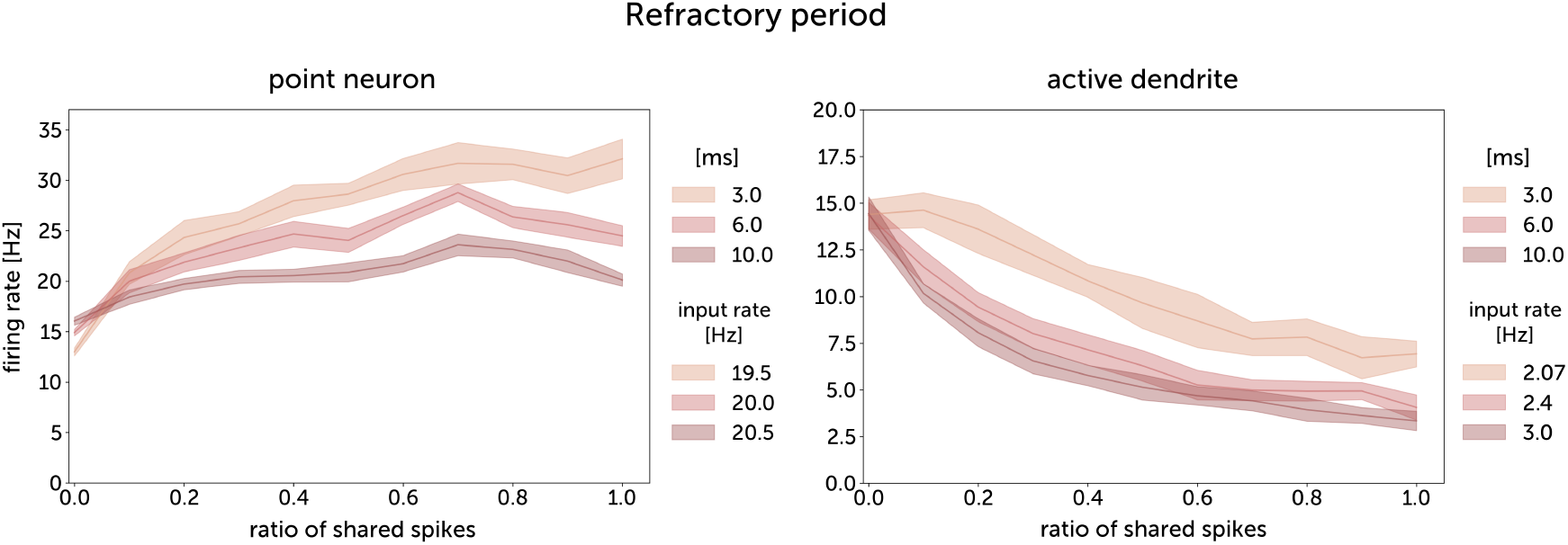
Somatic firing rate responses to correlated synaptic activity for an integrate-and-fire point neuron and for an integrate-and-fire neuron with dendrite. The synaptic conductances were 0.5 nS for a neuron with a dendrite, and 0.105 nS for a point neuron. The time constant of jitter for the generation of correlated input spike trains, *τ_j_* was 10 ms.

### 2.3 Simplified discrete-state dendritic model

The simple space-time illustration of collisions of dendritic spikes is given in Fig. 7a. We can distinguish two modes of interference between spikes: 1) direct collisions 2) inability to initiate/propagate spike due to refractory period. Assuming a constant velocity of propagation of the dendritic spikes and a uniform distribution of dendritic spikes, the number of direct collisions is proportional to the length of the dendrite, while the sole effect of refractoriness doesn’t scale with the length of a dendrite. Finally, we designed a mathematical model, called *discrete-state model*, aiming at grasping the core mechanism of spike propagation / annihilation. Compared to the more detailed models described so far, this one has much smaller simulation time and allows to easily modify parameters independently (*e.g.* propagation speed). The simulation is exact in that it is not based on a (time) discretization of the propagation of dendritic fronts. Finally, it allows to see whether the annihilation of spikes is enough to explain decrease of mean somatic spikes number. More precisely, suppose that we are given a set of spatio-temporal synaptic inputs 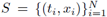 and assume that each input (*t_i_, x_i_*) produces two contra-propagating fronts (*e.g.* dendritic spikes). We also assume that *S* is generated with the procedure described above. In Figure 7b, the synaptic inputs are the yellow dots and the lines, the propagating fronts. Starting from *S*, we build the set of fronts and annihilation events recursively (see Methods). Figure 7b shows the networks of propagating fronts for different propagating speeds but same inputs *S*. One can see that increasing speed increases the dendritic spikes reaching the soma (*x* = 0). Intuitively, this occurs because a front has less chance to meet another front for higher propagation speeds. Finally, in Fig. 7c we present the dependency of the mean number of spikes reaching the soma as a function of the ratio of shared spikes. In this scenario, each synaptic input triggers a dendritic spikes, so the results corresponds to the case of large synaptic weights of the multi-compartment Hodgkin-Huxley model (Fig. 4c).

**Fig.7.**
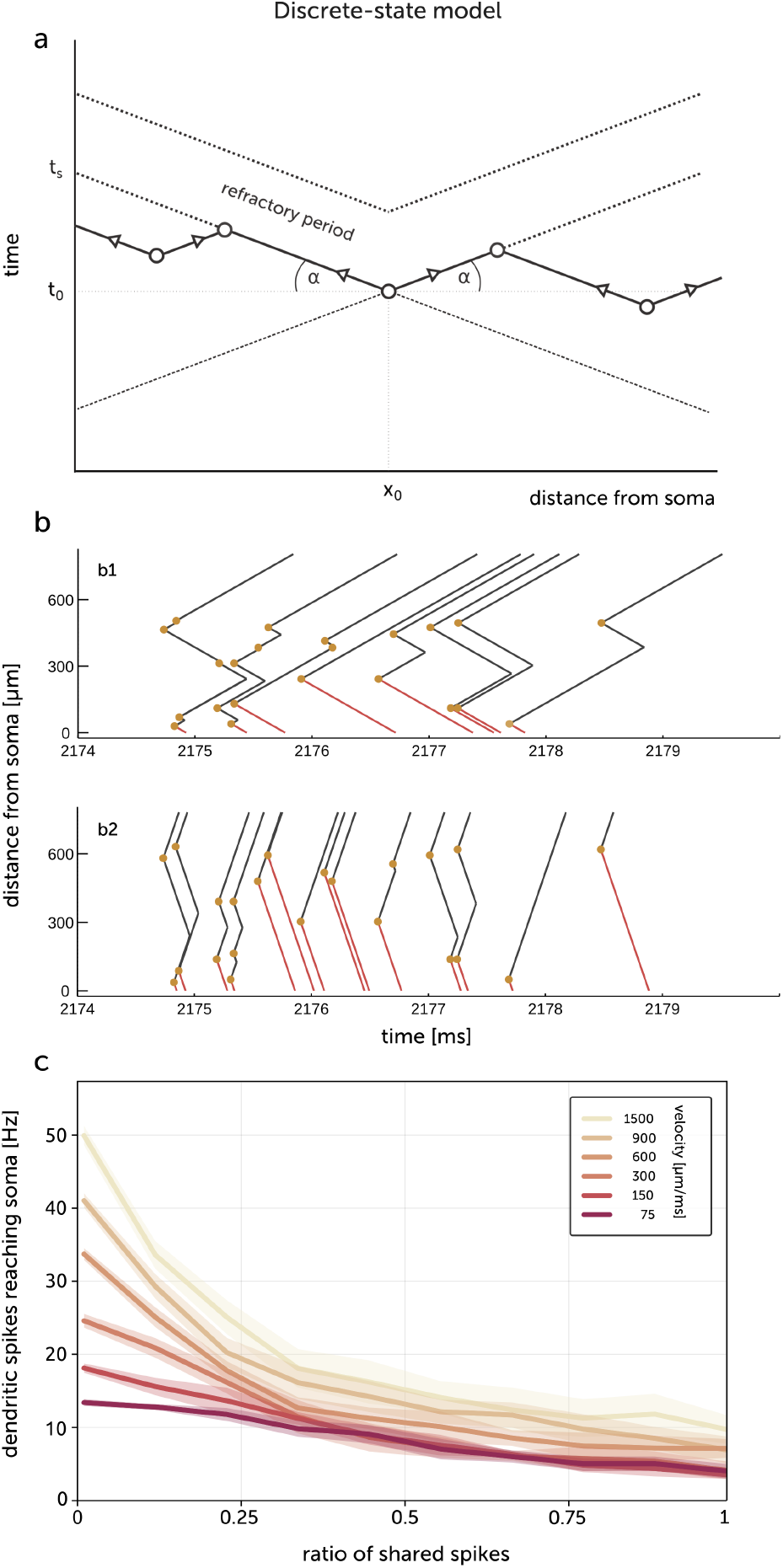
Discrete state dendritic model. **(a)** Illustration of propagations and collisions of dendritic spikes. The dendritic spike is created at point (*t*_0_, *x*_0_) and propagates to the soma and to the distal end of a dendrite. Spikes created in the area enclosed between the wavefronts and their extensions can collide with the spike created at point (*t*_0_, *x*_0_). Moreover, spikes cannot be created in the refractory area. The velocity of the spike is 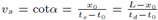, where *L* is a length of a dendrite and *t_s_* and *t_d_* are the times when spike reaches soma and distal end of a dendrite respectively. The spikes created in space-time area 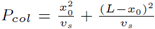 can collide with the spike created at point (*t*_0_, *x*_0_). The space-time area in a refractory state is *P_ref_* = *Lt_r_*. Assuming that a density of probability of spike creation *p_s_*(*t*, *x*) is uniform and assuming normalization 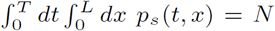, where *N* is a total number of spikes created in time *T*, the number of collisions should be proportional to *p_s_P_col_* and increases linearly with the length of the dendrite *L*, while the number of spikes not created due to refractory period *p_s_P_ref_* doesn’t scale with *L*. **(b)** Propagations of spikes in a discrete-state dendritic model. The origins of spikes are marked by yellow circles. Red lines indicates dendritic spikes which reach the soma. The velocity of dendritic spikes is 4 times higher in (b2) than in (b1). The ratio of shared input spikes, *c_G_* = 0.5. **(c)** Dependence of the mean rate of dendritic spikes reaching the soma on global correlation for the discrete-state model for different velocities of the dendritic spikes. Time constant of the jitter *τ_j_* = 2 ms.

## 3 Discussion

In this paper, we have investigated the behavior of excitable neuronal dendrites using different models. The main property that we explored is the ability of the dendrite to initiate and propagate fast dendritic spikes and its consequences on processing correlated synaptic activity.

One of our main findings is that the presence of fast dendritic spikes endow neurons to process inputs according to a mode of inverse correlation processing, in which the neuron fires less when the inputs are correlated. This is contrary to point neurons, which generally show the opposite mode, where firing is directly increased by correlations [47–50]. However, point neurons can display inverse correlation processing only for very long refractoriness or for extreme stimulation conditions.

We have further shown that this inverse correlation processing mode was present in different models, such as a Hodgkin-Huxley model of fast sodium dendritic spikes, an integrate-and-fire model for dendrites, or a discrete-state dendritic model. We discuss below possible implications of this finding, and ways to test them experimentally.

Our model challenges the well-known finding that the presence of correlations in the pre-synaptic activity necessarily augments the firing of neurons [47–50]. We find here that neurons equipped with excitable dendrites can operate in a regime which is opposite: neurons with active dendrites generally decrease their firing when correlations are present in their afferent activity. Furthermore, this inverse correlation processing mode was dominant in models where dendrites generate fast spikes (such as Na^+^ spikes) whenever the intensity of synaptic bombardment was high enough. Recent experiments [44] provided strong evidence that Na^+^ dendritic spikes are much more frequent than somatic spikes in awake animals (10 times more frequent during exploration), so there is a serious possibility that neurons can be set to this inverse correlation processing mode.

It is interesting to ask how the response to synaptic input correlation can be affected by the morphology of dendritic tree. In our linear model of dendrite, the length of the distal dendritic part is much shorter than the total length of distal branches present in more complex morphologies. The initiation of sodium spike is enhanced in more distal branches where higher input resistance causes higher local depolarization triggered by a single synaptic input. The resulting increase of the spike rate in distal dendrites can lead to the higher rate of spike annihilations and therefore amplified inverse correlation processing. Indeed, the collisions of dendritic spikes can effectively prevent the propagation of sodium spikes from distal branches to the soma [56].

While it is beyond the scope of the present study, one can ask if NMDA and Ca^2+^ spikes may have similar impact on correlation processing. The NMDA spike, however, is different than the Na^+^ spike because it cannot usually propagate (it would require continuous glutamatergic stimulation along the dendrite). Therefore interactions between NMDA spikes cannot be as global as interactions between Na^+^ spikes. Calcium spikes on the other hand propagate usually in a restricted area of the apical dendritic tree and cannot actively invade soma [57], while their frequencies are much lower than frequencies of fast sodium spikes [44, 58].

We demonstrated the inverse correlation phenomenon also on a simplified integrate-and-fire-type model, which can be extended to include neuronal adaptation (AdEx model). Although this type of model is usually applied to point neurons without any spatial extent, we extended it over multiple compartments connected through axial currents such that propagating spikes can be simulated. The AdEx model allows to simulate different intrinsic properties such as bursting and adaptation, [55,59]. This allows us to later extend the present model by including such mechanisms in dendrites. The AdEx model is also compatible with neuromorphic hardware, [60], and which is presently extended to include neurons with excitable dendrites [61]. In addition, to allow for analytical treatment of the dendritic spikes, we further simplified the model to a discrete state model capturing the dynamics of the collisions of dendritic spikes.

What are the consequences of such inverse correlation processing? One obvious effect will be to cancel correlations at the network level, because the neurons subject to correlated input will fire less, and neurons with decorrelated input will dominate the population, and will most likely shift the population towards decorrelated firing. Such dynamics should be examined by future network models with neurons equipped with excitable dendrites.

Finally, it is important to test the inverse correlation processing experimentally. Dynamic-clamp was used previously to investigate the fact that neurons increase their synchrony [62], but unfortunately, this technique cannot be used to test the present mechanism, because dynamic-clamp emulates inputs in the soma (the site of the recording electrode), while it is important here that the inputs are dendritic. One possibility would be to use 2-photon imaging with glutamate uncaging [63] to simulate inputs with controlled frequency and synchrony. Another possibility would be to use voltage-sensitive dye imaging of dendrites *in vitro* [64] combined with controlled network activity in the slice, to directly monitor the genesis, propagation and possible collision of dendritic spikes.

## 4 Methods

### 4.1 Hodgkin-Huxley dendritic model

We used a model in which somatic compartment was connected with a linear dendrite. The morphological and electrophysiological parameters were chosen to obtain the realistic membrane depolarization at soma and local membrane depolarization as a function of distance of synapse from soma [65], [see Appendix, Fig 8]. The diameter of a soma was 40 *μm* and the diameter of the dendrite was gradually decreasing with the distance from soma. In most of simulations there were 200 dendritic compartments of the length 5 *μ*m; for Fig. 5c there were 50 dendritic compartments of length 20 *μ*m.

**Fig.8.**
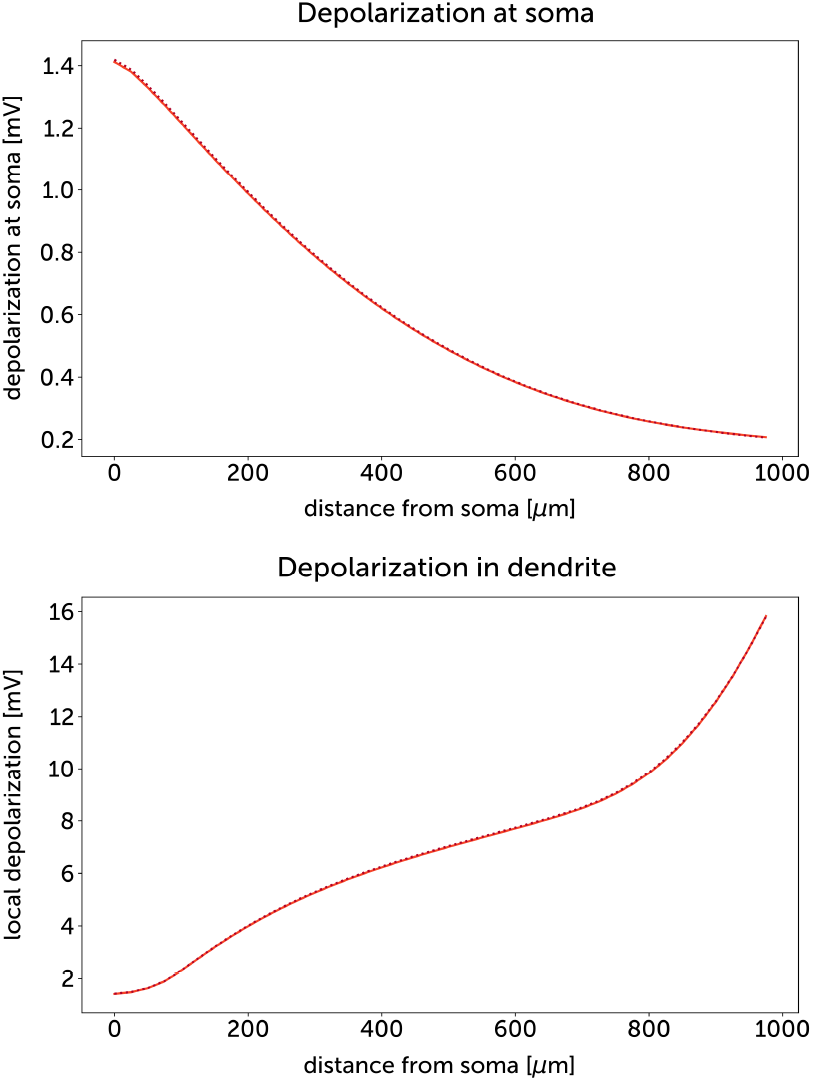
Depolarization in soma and dendrites following synaptic input. *Top:* The depolarization in soma for synaptic inputs at different dendritic locations. *Bottom:* The local dendritic depolarization is shown in the same conditions. The synaptic conductance was 0.5 nS.

The voltage-gated fast sodium and delayed-rectifier potassium ion channels were present in soma and in dendrite. The cable equation was as following:

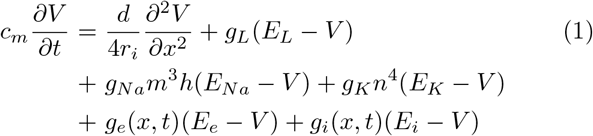

where *c_m_* is a specific capacitance, *d* is a diameter of a dendrite, *r_i_* is an intracellular specific resistance. *g_L_* is a leak conductance, *E_L_* is a leak reversal potential, *g_Na_* is a maximal conductance of fast sodium channels, *E_Na_* is a reversal potential of sodium current, *g_K_* is a maximal conductance of potassium current, *E_K_* is a reversal potential of a potassium current. *g_e_* (*g_i_*) is a conductance of excitatory (inhibitory) synapses, *E_e_* (*E_i_*) is a reversal potential of excitatory (inhibitory) synapses. *m*, *h* and *n* represent gate states of ion channels. The gate states evolves according to equations:

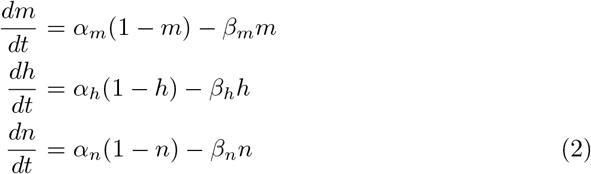

Where

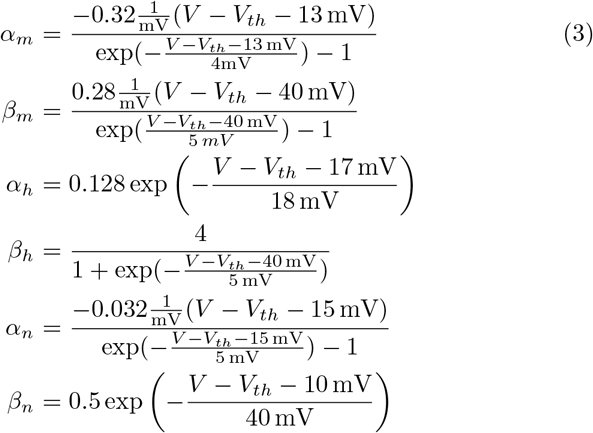

Whenever there is excitatory (e) or inhibitory (i) synaptic event: *g_s_*(*t*) → *g_s_*(*t*) + *δg_s_*(*t*). The value of *δg_s_*(*t*) is a weight of a synapse. Between events synaptic conductances follow equation:

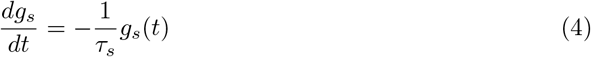

where *τ_s_* is a time constant of synaptic current.

The number of simultaneous synaptic events needed to initiate dendritic sodium spike depends on the distance between synapse and soma, [see Appendix, Fig 9]. Few synaptic events are needed to initiate sodium spike in the distal tip of the dendrite. This aspect of dendritic sodium spike initiation was studied in more detailed computational models [43].

**Fig.9.**
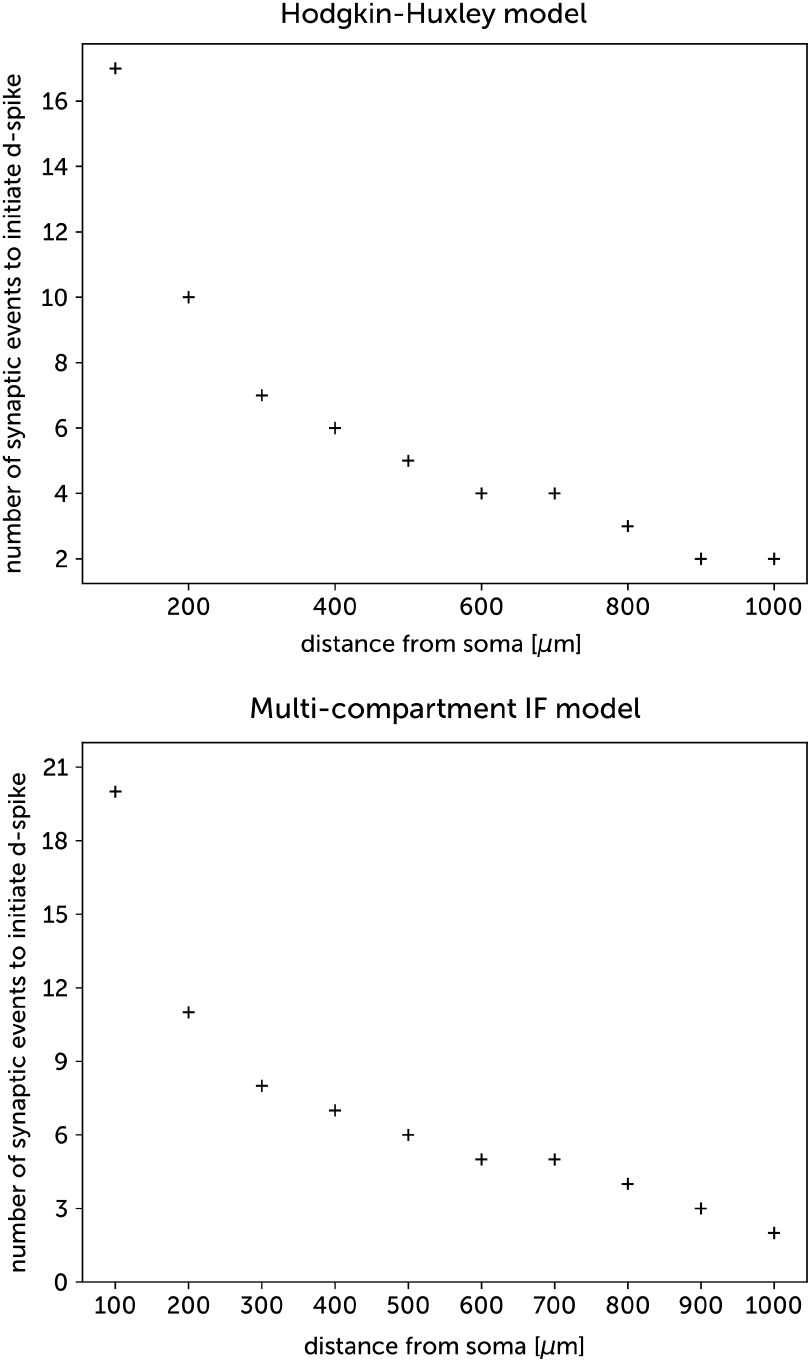
Number of synaptic events needed to initiate a dendritic spike in a Hodgkin-Huxley model and in a multi-compartment integrate and fire model. The synaptic conductance was equal to 0.5 nS for both models.

The velocity of dendritic sodium spikes decreases with the distance from soma starting with 1200 *μ*m/ms in the proximal dendrite to 200 *μ*m/ms in the distal dendritic end (Fig. 10), similarly to experimental observations [66].

**Fig.10.**
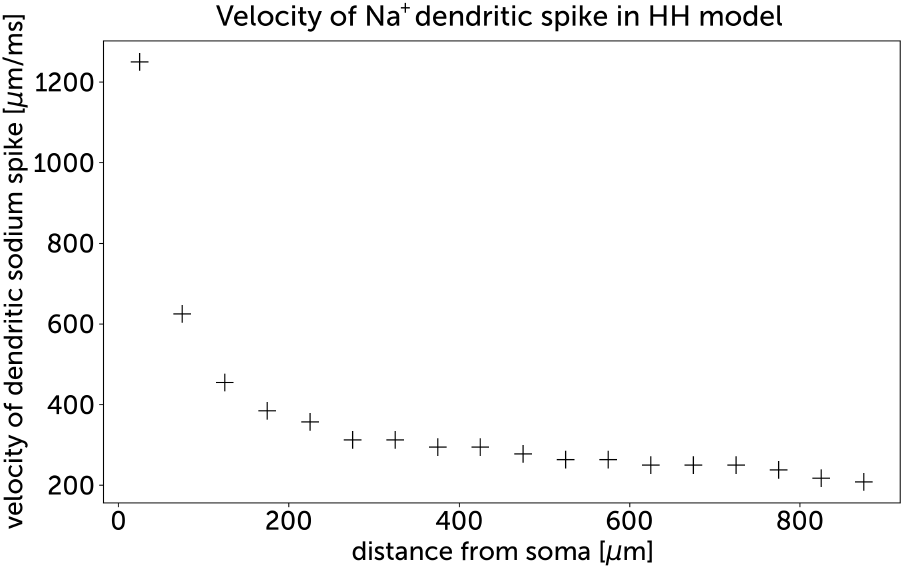
Velocity of dendritic sodium spikes as a function of the distance from the soma in a Hodgkin-Huxley model.

In our simulations: *c_m_* = 1 *μ*F*/*cm^2^, *r_i_* = 100 *Ω*cm, *d* = 1 *μ*m, *g_L_* = 100 *μ*S*/*cm^2^, *E_L_* = −70 mV, *g_Na_* = 12 mS/cm^2^, *E_Na_* = 58 mV, *gK* = 7 mS=cm^2^, EK = –80 mV, *V_th_* = –63 mV, *Ee* = 0 mV, *E_i_* = –75 mV, *τ_s_* = 5 ms.

### 4.2 Integrate and fire dendritic model

We used exponential integrate and fire model, which can be easily extended by neural adaptation mechanism, forming AdEx model [55]. In exponential integrate and fire model a spike is triggered by the exponential depolarizing current [67] which provides smooth spike initiation zone in place of fixed threshold of Leaky Integrate and Fire (LIF) model. For a multi-compartment model a membrane voltage *V* (*t*, *x*) is governed by a following cable equation:

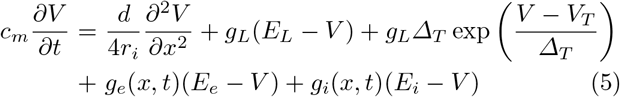

where *V_T_* is a spike threshold and *∆_T_* is a slope factor.

This equation is solved numerically by discretization of the equations over multiple dendritic compartments. When a membrane potential of any compartment is near threshold *V_T_* depolarizing exponential current surpass other currents and membrane voltage quickly tends to infinity. Whenever the membrane potential crosses peak value *V_p_* we detect a spike and compartment enters into refractory period in which voltage is repolarized according to equation:

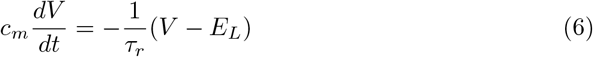

The exponential decay characteristic of this equation models the falling phase of an action potential. The time constant of repolarization *τ_r_* was chosen in such a way that after 1 ms the difference between *V* and *E_L_* was smaller than 0.01 mV. After emitting a spike the compartment stays in the refractory state during time *t_r_*.

The number of simultaneous synaptic events needed to initiate dendritic spike depends on the distance between synapse and soma, [see Appendix, Fig 9]

In our simulations: *c_m_* = 1 *μ*F*/*cm^2^, *r_i_* = 100 *Ω*cm, *d* = 1 *μ*m, *g_L_* = 100 *μ*S*/*cm^2^, *E_L_* = −70 mV, *V_T_* = −50 mV, *V_p_* = −20 mV, *τ_r_* = 117 *μs/c_m_*, *∆_T_* = 2 mV, *E_e_* = 0 mV, *E_i_* = −75 mV, *τ_s_* = 5 ms.

### 4.3 Simulation of the discrete-state model

Given a set S of somatic inputs, we describe our procedure to re-construct the propagating dendritic fronts leading to somatic spikes. Take the latest element *s*_1_ = (*t*_1_, *x*_1_) ∈ *S*. We look for possible annihilation events with its outgoing front, *i.e.* going away from the soma located at *x* = 0. Hence, we select all inputs *S^↑^*(*s*_1_) in *S* \ {*s*_1_} which are located in the cone above *s*_1_ defined as *S^↑^*(*s*_1_) *≡* {(*t*, *x*): *x* ≥ ±(*t* − *t*_1_) + *x*_1_}. Consider the somatic input *s*_2_ ∈ *S^↑^*(*s*_1_) closest to the line *x* = −(*t* − *t*_1_) + *x*_1_: it will produce an annihilation event with *s*_1_. Perform the same for *s*_2_ recursively until possible. This gives a chain of annihilation events (*s*_1_, *s*_2_, …, *s_k_*). We do the same for possible annihilation events with the ongoing front from *s*_1_, *i.e.* going toward the soma located at *x* = 0. It gives another chain of events (*s_−p_*, …, *s*_0_, *s*_1_). Hence, we found a chain of annihilation events *C*(*s*_1_) = (*s_−p_*, …, *s_k_*): it produces **exactly one** somatic spike. We continue the same procedure with *S*\*C*(*s*_1_) until *S* =∅. The overall procedure runs in *O*(*n*^2^) where *n* is the cardinal of *S*.

### 4.4 Correlation measure for spike trains

To measure the correlation between two spike trains *S_i_*(*t*) and *S_j_*(*t*) we applied the cross-covariance function CCVF_*ij*_(*s*) = ⟨*S_i_*(*t*)*S_j_*(*t* + *s*) ⟩ − ⟨*S_i_*(*t*) ⟩ ⟨*S_j_*(*t*) ⟩, where ⟨*…* ⟩ denotes a time average. The normalized integral 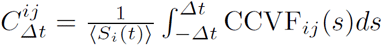 tells us about a correlation within a time window[–*Δt*, *Δt*] [Fig. 1]

## Acknowledgements

We would like to thank Gilles Wainrib for fruitful discussions. Supported by CNRS, INRIA and the European Community (Human Brain Project, FP7-604102 and H2020-720270). All simulations were done using the Brian 2 neuronal simulator and Python 3 language. The code can be downloaded from the ModelDB repository: http://modeldb.yale.edu/244700.

## Appendix

In this appendix, we show additional properties related to the dendritic models. The profile of depolarization, in soma and dendrite, caused by a synaptic input is shown in Fig 8. The numbers of synaptic inputs needed to initiate dendritic spike are shown in Fig 9, and the velocity of sodium dendritic spike in Fig 10.

